# The glucocorticoid dexamethasone inhibits HIF-1alpha stabilisation and metabolic reprogramming in lipopolysaccharide-stimulated primary macrophages

**DOI:** 10.1101/2023.09.20.558626

**Authors:** Sally A Clayton, Chloe Lockwood, John D O’Neil, Kalbinder K Daley, Sofia Hain, Dina Abdelmottaleb, Oliwia O Bolimowska, Daniel A Tennant, Andrew R Clark

## Abstract

Synthetic glucocorticoids are used to treat many chronic and acute inflammatory conditions. Frequent adverse effects of prolonged exposure to glucocorticoids include disturbances of glucose homeostasis, caused by changes of glucose traffic and metabolism in muscle, liver and adipose tissues. Macrophages are important targets for the anti-inflammatory actions of glucocorticoids. These cells rely on aerobic glycolysis to support various pro-inflammatory and antimicrobial functions. Employing a potent pro-inflammatory stimulus in two commonly-used model systems (mouse bone marrow-derived and human monocyte-derived macrophages), we showed that the synthetic glucocorticoid dexamethasone inhibited lipopolysaccharide-mediated activation of the hypoxia-inducible transcription factor HIF-1α, a critical driver of glycolysis. In both cell types, dexamethasone-mediated inhibition of HIF-1α reduced the expression of the glucose transporter GLUT1, which imports glucose to fuel aerobic glycolysis. Aside from this conserved response, other metabolic effects of lipopolysaccharide and dexamethasone differed between human and mouse macrophages. These findings suggest that glucocorticoids exert anti-inflammatory effects by impairing HIF-1α-dependent glucose uptake in activated macrophages. Furthermore, harmful and beneficial (anti-inflammatory) effects of glucocorticoids may have a shared mechanistic basis, depending on alteration of glucose utilisation.

**GRAPHICAL ABSTRACT:** 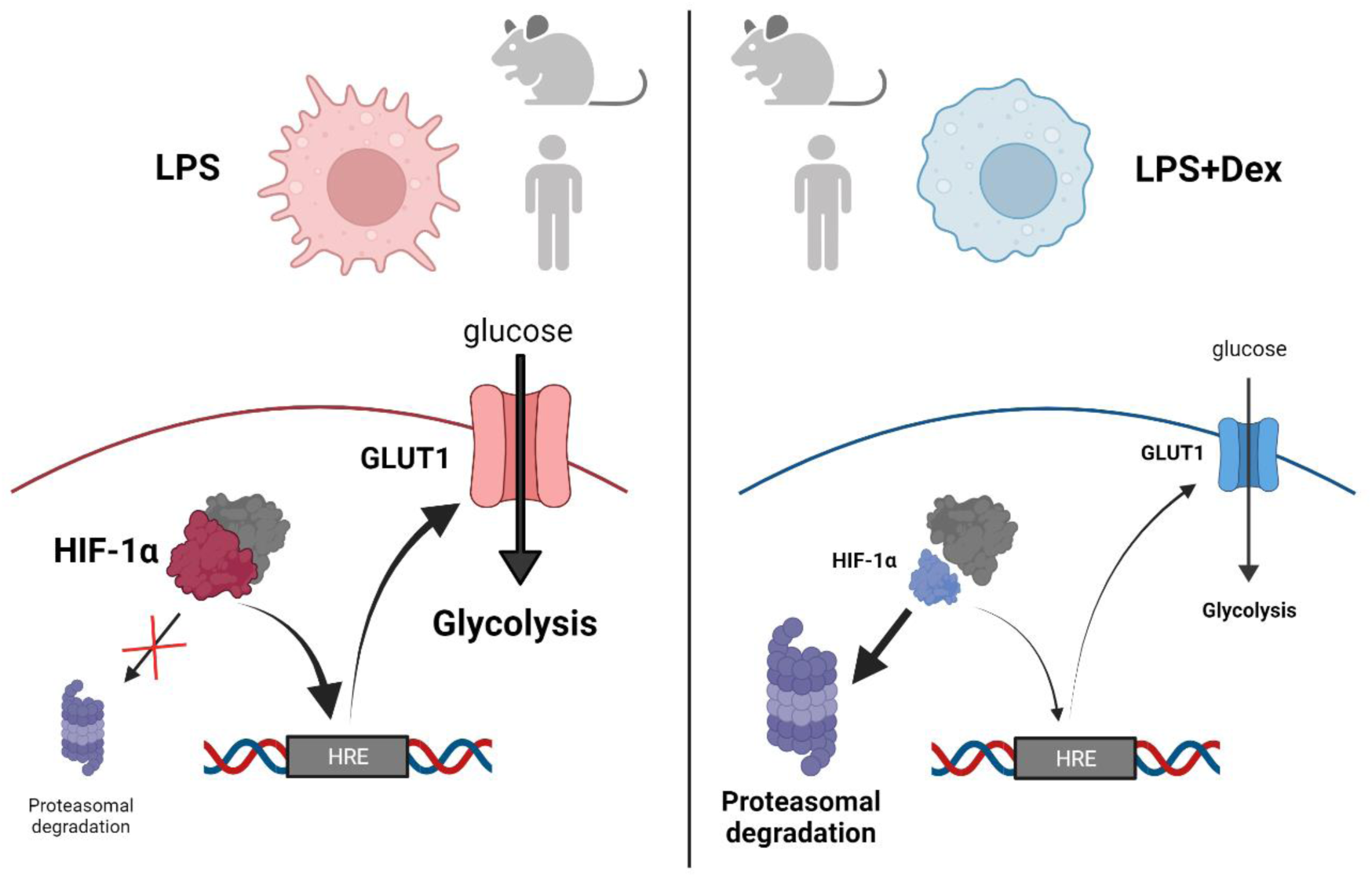

## INTRODUCTION

Glucocorticoids (GCs) are steroid hormones that are mainly produced by the adrenal glands, and fulfil important roles in adaptive physiological responses to stress. One of their most important functions is to maintain the supply of glucose to the brain, an extremely glucose-dependent organ that constitutes around 2% of human body mass but consumes around 50% of available glucose (1, 2). The maintenance of glucose availability is achieved by modulating glucose homeostatic processes in metabolic organs such as liver, muscle, adipose and pancreas; preventing glucose uptake and utilisation, promoting glucose synthesis (gluconeogenesis) and mobilisation (3, 4). Profound anti-inflammatory and immuno-suppressive effects of the endogenous GC cortisone were discovered in the mid twentieth century by Hench and colleagues (5). Since then, synthetic GCs (highly similar in structure to cortisone) have been very extensively used to treat chronic and acute immune-mediated inflammatory diseases. Most recently, the GC dexamethasone was found to reduce mortality in patients infected with the zoonotic virus SARS-CoV-2 and requiring respiratory support due to lung damage caused by the virus-induced cytokine storm (6).

Cumulative exposure to synthetic GCs is associated with adverse effects in several organs and tissues (7). Sometimes severe, even life-threatening, these resemble the consequences of endogenous GC excess, known as Cushing’s syndrome. Many (perhaps most) harmful effects of synthetic GCs are clearly related to the normal functions of endogenous GCs in regulating systemic metabolism. It is misleading to label such harmful responses to drug as “side effects”, since this term implies an unpredictable, off-target phenomenon (8). The anti-inflammatory effect of GCs at first seems a maladaptive response to stress, but is now also rationalised as a means of conserving glucose for the brain by reducing its utilisation in peripheral tissues. This is because activated immune cells often increase glycolytic metabolism, which is accompanied by rapid uptake and consumption of glucose (9). Classically associated with low oxygen environments such as poorly vascularised tissues or tumors, glycolysis occurs in activated immune cells regardless of oxygen availability (hence known as aerobic glycolysis). It permits very rapid increases in biosynthetic and other energy-requiring processes. For example glycolysis is required for phagocytosis, migration, killing of intracellular pathogens and production of inflammatory cytokines by activated macrophages (10–16).

Increased glucose consumption at sites of inflammation has been known for decades (17) but its molecular basis is only now becoming understood, with the rapid growth of the field of immunometabolism (18). Much of what we know in this domain is based on studies of mouse macrophages. Important metabolic differences between human and mouse macrophages have been described, but are often under-appreciated (18–22). Species differences are important to consider when translating findings from experimental models of immune-mediated inflammatory disease, attempting to discover new immunometabolism-targeting treatments or improve the safety profile of existing drugs (23).

As in hypoxic tissues, the glycolytic program of activated immune cells is orchestrated by the hypoxia-inducible transcription factor HIF-1, a dimer consisting of a hypoxia-sensitive subunit (HIF-1α) and a constitutively-expressed partner (HIF-1β). In normoxia HIF-1α is constitutively synthesised, hydroxylated by oxygen-dependent prolyl hydroxylases (PHDs), then recognised and ubiquitinated by the von Hippel Lindau ubiquitin ligase complex, and subsequently degraded by the proteasome. Depletion of oxygen prevents the hydroxylation step, leading to stabilisation and accumulation of HIF-1α, which then combines with HIF-1β to bind to hypoxia response elements (HREs) and drive expression of glycolytic genes and others involved in cellular adaptation to hypoxia. In mouse myeloid cells stimulated with lipopolysaccharide (LPS), disruption of the mitochondrial tricarboxylic acid (TCA) cycle leads to accumulation of succinate, which inhibits prolyl hydroxylases and promotes HIF-1α accumulation independently of oxygen depletion (13, 24). HIF-1α-mediated expression of *Nos2* (nitric oxide synthase 2, or inducible nitric oxide synthase, iNOS) (14, 25, 26) causes production of nitric oxide, which impairs the function of mitochondrial electron transport chains and enzymes of the TCA cycle, reducing the efficiency of ATP production by oxidative phosphorylation and reinforcing the cell’s dependence on glycolysis (26–28). HIF-1α binds to and activates the *Il1b* promoter, ultimately leading to production of the pro-inflammatory cytokine IL-1β (13, 14, 29). HIF- 1α-mediated regulation of other pro-inflammatory mediators has been inferred but in most cases not yet directly demonstrated (13). According to this model the mechanisms of activation of HIF-1α under hypoxia and LPS stimulation are fundamentally similar, depending on changes of prolyl hydroxylase activity.

Here we hypothesised that GCs act upon immune cells in much the same way that they act upon metabolic tissues, restricting utilisation of glucose as a means of conserving it for the brain under conditions of stress. To test this hypothesis we used a variety of metabolic and gene expression assays in both human and mouse primary macrophages, in order to highlight both commonalities and differences.

## MATERIALS AND METHODS

### Macrophage isolation and culture

Blood from anonymous healthy donors was obtained in the form of leukapheresis cones from the NHS Blood and Transplant Service (ethical approval ERN_16-0191). Monocytes were isolated by negative selection using RosetteSep Human Monocyte Enrichment Cocktail (STEMCELL #15068) and Ficoll density gradient centrifugation, and were differentiated into macrophages by culture in RPMI 1640 + L-glutamine (Gibco #21875034) supplemented with 5% heat-inactivated FCS (LabTech #80837) and 50 ng/ml M-CSF (PeproTech #300-25) for 7 days.

Wild type C57BL/6J mice were housed at the University of Birmingham Biomedical Services unit, and all maintenance and procedures were carried out according to the Home Office guidelines and approved by the University of Birmingham Animal Welfare and Ethical Review Board. Mice were sacrificed by cervical dislocation and legs removed for bone marrow isolation. Femurs and tibiae were cleaned of muscle tissue and cut at each end. Bone marrow was extracted by centrifugation and plated in RPMI 1640 + L-glutamine (Gibco #21875034) supplemented with 10% heat-inactivated FCS (Sigma Aldrich #F2442) and 50 ng/ml M-CSF (PeproTech #300-25) for 7 days to differentiate bone marrow-derived macrophages (BMDMs).

For stimulations, cells were seeded at 0.5x10^6^ cells/ml (human) or 1x10^6^ cells/ml (mouse) in tissue culture-treated 12-well or 6-well plates unless otherwise stated. Stimulation medium was prepared with LPS (E. coli, Serotype EH100 (Ra) (TLRgrade™) – Enzo Life Sciences #ALX-581-010-L002) at 10 ng/ml and/or dexamethasone dissolved in DMSO (Sigma Aldrich # D8893) at 100 nM, without M-CSF. DMSO concentration was matched across all conditions.

### Seahorse Metabolic Flux Assays

Cells were seeded at 50,000 cells/well (human) or 80,000 cells/well (mouse) in Agilent Seahorse XFe96 cell culture microplates and left overnight to adhere. A combined version of the standard Mito and Glyco stress tests was carried out as described (30). Seahorse XF RPMI medium, pH 7.4 (Agilent #103576-100) was supplemented with 2 mM L-glutamine (Sigma Aldrich #G7513). The following injection protocol was used (final assay concentrations): (A) D-glucose (10 mM), (B) oligomycin A (human, 1 μM; mouse, 1.5 μM), (C) carbonyl cyanide p-trifluoromethoxyphenylhydrazone (FCCP) (human, 5 μM; mouse, 1.5 μM) + sodium pyruvate (1 mM), and (D) rotenone (100 nM) + antimycin A (1 μM) + 2-deoxy-D-glucose (20 mM). Calculations of metabolic parameters were carried out according to recommendations by Seahorse (Agilent).

For ATP production rate assays, Seahorse XF RPMI medium, pH 7.4, was supplemented with 2 mM L-glutamine, 10 mM D-glucose (Sigma Aldrich #G7021) and 1 mM sodium pyruvate (Sigma Aldrich #P5280). The following injection protocol was used: (A) oligomycin A (human, 1 μM; mouse, 1.5 μM), (B) rotenone (100 nM) + antimycin A (1 μM). Analysis was performed using XF Real-Time ATP Rate Assay Report Generator (Agilent).

For acute stimulation assay BMDMs were seeded as above except at 100,000 cells/well. At the start of the assay D-glucose was injected to final concentration of 10 mM, with vehicle, LPS (10 ng/ml), Dex (100 nM) or both. ECAR was measured every 5 minutes for 4.5 h. At the end of this time 2- deoxy-D-glucose was added and ECAR measured every 5 minutes for a further 15 minutes to confirm that measured ECAR was dependent on glycolysis. Change in ECAR was calculated relative to the start of assay.

Drugs were purchased from Sigma Aldrich (2-deoxy-D-glucose #D8375; antimycin A #A8674) or Cayman Chemical Company (oligomycin A #11342; FCCP #15218; rotenone #83-79-4). Assay normalisation was carried out by calculating a viable cell count ratio. Immediately following assay completion, cells were incubated with calcein-AM viability dye (eBioscience, 65-0853-78) at 1 μM in phosphate-buffered saline (PBS) for 30 min at 37°C. Fluorescence was measured by plate reader (excitation, 490 nm; emission, 515 nm).

### Lactate/glucose measurements

Lactate and glucose concentrations in conditioned medium from macrophage cell cultures were measured using the Nova Stat Profile Prime cell culture analyser. Glucose consumption was calculated by subtracting concentration in normal culture media (RPMI 1640) from values obtained from conditioned medium samples.

### RT-qPCR

RNA was isolated using Norgen Total RNA Purification Plus kit (Geneflow, P4-0016) according to manufacturer’s protocol, and quantified by Nanodrop (Thermo Fisher). cDNA was synthesised using iScript Reverse Transcriptase (Bio-Rad #1708891) from 250 ng RNA/reaction. Gene expression was measured by reverse transcription quantitative polymerase chain reaction (RT-qPCR) using the Bio-Rad CFX384 system. SYBR TB Green Premix Ex Taq (Takara, #RR820W) and primers supplied by Sigma Aldrich (see table 1) were used unless otherwise stated. *UBC* (human) or *Rpl13a* (mouse) were used to normalize mRNA measurements via 2^−ΔΔCt^ method.

**Table 1:**
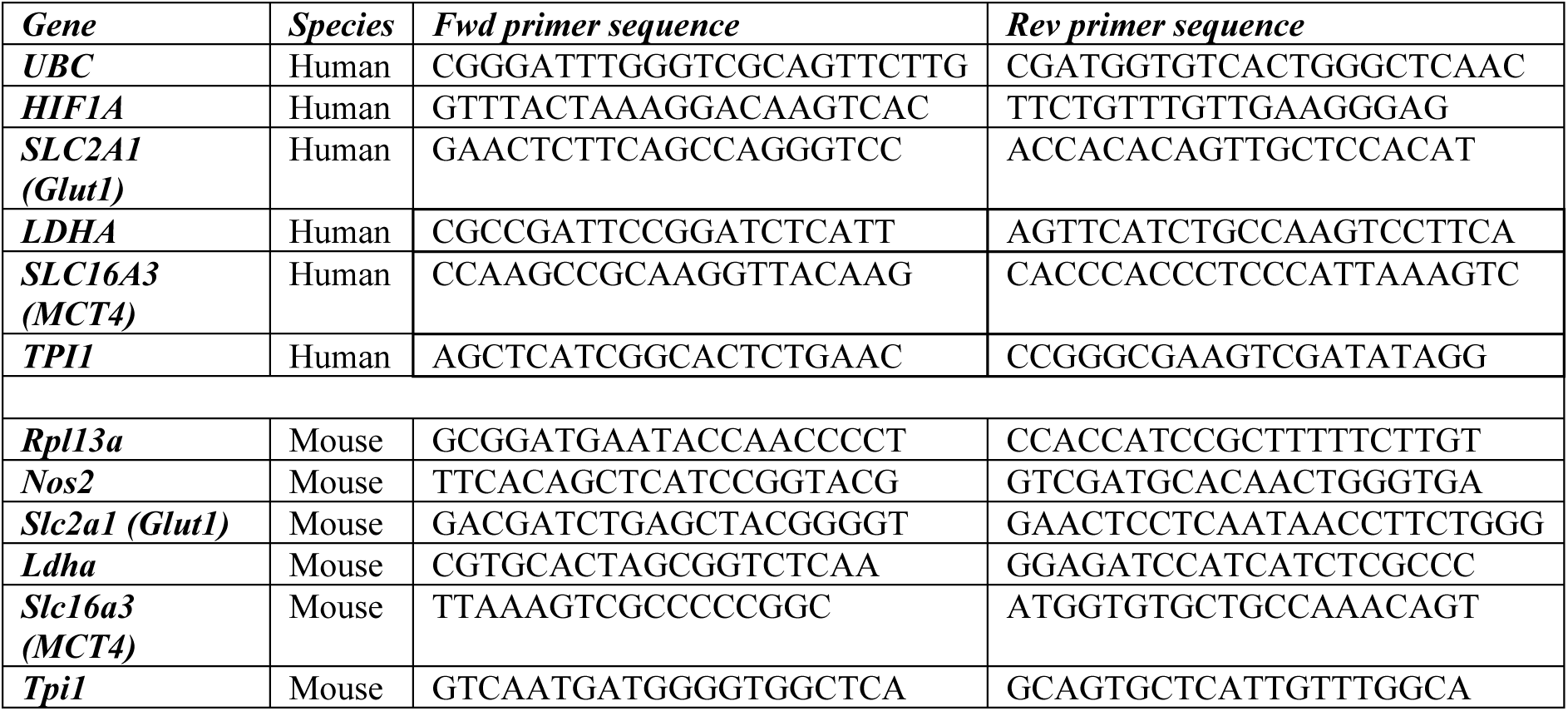
RT-qPCR primer sequences for human and mouse.

Mouse *Hif1a* gene expression was measured using ThermoFisher TaqMan gene expression assay (assay ID: Mm00468869_m1) normalised to Rpl13a (assay ID: Mm05910660_g1), using Applied Biosystems TaqMan Gene Expression Master Mix (#4369016).

### Western blotting

Cells were lysed in radioimmunoprecipitation assay (RIPA) buffer or directly into 2X XT Sample buffer (Bio-Rad #1610791) + 1X XT Reducing Agent (Bio-Rad #1610792) (for detecting human HIF-1α protein). Samples were passed through a QIAshredder column to remove genomic DNA (QIAGEN, 79656). If lysed in RIPA buffer, protein was quantified by Pierce BCA Protein Assay (ThermoFisher #23225), XT sample buffer + reducing agent, or Laemmli loading buffer + β- mercaptomethanol was added (to 1X), and an equal protein mass loaded onto the gel. If lysed in Sample buffer, an equal volume of each sample was loaded. Before loading, samples were heated to 95°C for 5 min (unless blotting for GLUT1). Western blotting was performed using XT Bis-Tris protein gels (Bio-Rad) and XT MES running buffer (Bio-Rad, 1610789), or Tris-glycine gels (Bio-Rad) and Tris-glycine SDS running buffer (Geneflow # B9-0034). Protein was transferred to Bio-Rad Trans-Blot polyvinylidene difluoride (PVDF) membranes (Bio-Rad, 1704157) using Bio-Rad Trans-Blot Turbo transfer system. Primary antibodies were applied overnight at 4°C as specified in Table 2. HRP-conjugated secondary antibodies were obtained from Cell Signaling Technologies (#7074, #7076) and applied for 1hour at room temperature. Imaging was performed using Clarity Enhanced Chemiluminescence substrate (Bio-Rad #1705061) and a ChemiDoc MP Imaging System (Bio-Rad). Densitometry was performed using ImageJ Figi.

**Table 2:** Antibodies used for Western blotting.

| **Protein** | **Supplier** | **Cat #** | **Host Species** | **Species used for** | **Molecular Weight** | **Dilution** | **Medium** |
| --- | --- | --- | --- | --- | --- | --- | --- |
| HIF-1α | BD Biosciences | 610969 | Mouse | Human | 120kDa | 1/500 | 1% Milk |
| HIF-1α | Cell Signalling | 14179 | Rabbit | Mouse | 120kDa | 1/1000 | 5% BSA |
| (pro)-IL-1-β | Abcam | ab254360 | Rabbit | Human, Mouse | 30kDa | 1/1000 | 5% Milk |
| iNOS | Abcam | ab202417 | Rabbit | Mouse | 131kDa | 1/1000 | 5% Milk |
| GLUT1 | Abcam | EPR3915 | Rabbit | Human | 54kDa | 1/1000 | 5% Milk |
| β-actin | Sigma Aldrich | A1978 | Mouse | Human | 42kDa | 1/2000 | 5% Milk |
| α-Tubulin | Sigma Aldrich | T9026 | Mouse | Human, Mouse | 55kDa | 1/2000 | 5% Milk |

### NO measurements

Conditioned medium from BMDM stimulations was harvested at 24 hours. Nitric oxide analysis was conducted using the Griess Reagent Kit (ThermoFisher #G7921), following the manufacturer’s instructions. Standards made up with 1mM Nitrite Solution (concentration range 0µM-200µM), absorbance measured at 548nm.

### HRE-luciferase reporter assay

HeLa cells were seeded at 5x10^6^ cells per 10 cm dish. After 24 h, in OptiMEM medium (#31985062, Gibco), cells were co-transfected with 10µg TK-Renilla (#E2241, Promega) and 10µg HRE-luciferase (a gift from Navdeep Chandel, this plasmid contains three copies of a HRE from the *Pgk1* gene; Addgene plasmid #26731) in a 1.5:1 ratio with FuGene (#E2311, Promega). One day post transfection, cells were seeded into 96-well plates and treated with either DMSO (vehicle control), 20 µM or 40 µM KC7F2 and placed in Normoxia (∼20% O_2_) or Hypoxia (1% O_2_). 24 hours after stimulation, cells were PBS washed and lysed in 20 µl/well of Passive Lysis Buffer on a plate shaker for 15 minutes. The dual reporter luciferase assay (#E1910, Promega) was conducted following the manufacturer’s instructions and luminescence was detected on a BioTek Synergy HT plate reader (integration time: 0.1 sec).

### RNAseq analysis

Raw FASTQ data files (Illumina HiSeq 2500 single read) were downloaded from the Gene Expression Omnibus (GSE:131364). Quality control was carried out using FastQC v0.11.9, and TrimGalore v0.6.6 was used to trim adapter sequences and low quality reads (Phred < 20). The trimmed reads were aligned using STAR 2.7.2b-GCC-8.3.0. First, genome indexing was performed using GRCm38-mm10 mouse reference genome; then FASTQ files were aligned to this reference genome and sorted by coordinate. Rsubread v2.6.1 was used to generate gene counts from the aligned reads. Principal component analysis was performed to check the clustering of samples on a CPM-normalised and log-transformed count matrix. DEseq2 v1.38.3 was used to identify differentially expressed genes with Benjamini-Hochberg procedure for FDR adjustment of p values.

### Flow cytometry

Human macrophages were harvested from culture using PBS+5mM Ethylenediaminetetraacetic acid (EDTA; Sigma Aldrich #E7889) and cell lifters (Fisher Scientific #08-100-240). Cells were stained with eBioscience™ Fixable Viability Dye eFlour780 (Invitrogen #65-0865-14) at 1:1000 in PBS for 20 minutes. For GLUT1 antibody staining, cells were fixed and permeabilized with the BD Biosciences Cytofix/Cytoperm kit (#554722) according to manufacturer’s instructions. Cells were incubated with 0.05 µg/µl of Human Fc Block (BD Biosciences #564219) for 20 minutes prior to antibody staining with anti-GLUT1 (Abcam #EPR3915) or isotype control (Rabbit IgG, Abcam #EPR25A) at 1.58 μg/ml for 30 minutes. After washing, cells were stained with Goat Anti-Rabbit IgG H&L Alexa Fluor 488 secondary (Abcam #ab150077) at 2 μg/ml for 30 minutes. Cells were washed and re-suspended in FACS buffer for analysis on the BD Biosciences LSRFortessa. Data analysis was performed using FlowJo v10.

### Statistical Analysis

Statistical analysis was performed using GraphPad Prism. Statistical tests and corrections used are indicated in the figure legends. *P < 0.05, **P < 0.01, ***P < 0.001, and ****P < 0.0001. Sample numbers (n) specified in the figure legends indicate biological replicates.

## RESULTS

### Dexamethasone opposes metabolic reprogramming of LPS-treated mouse macrophages

As expected on the basis of previous reports (13, 22, 27, 28), 24 h treatment of mouse bone marrow-derived macrophages (BMDM) with LPS caused an increase in glycolysis (Fig. 1A, B) and a decrease in both basal and maximal respiration rates (Fig. 1C, D). This metabolic reprogramming was also demonstrated in separate ATP rate assays, which estimate the extent to which the generation of ATP is dependent on glycolysis or respiration (Fig. 1E). In the ATP rate plot, increased glycolysis is shown by a rightward shift and decreased OxPhos by a downward shift. LPS-induced changes in both glycolysis and OxPhos were opposed by simultaneous addition of a moderate dose of the synthetic glucocorticoid dexamethasone (Dex) (Fig. 1A-E). LPS also caused increases of glucose uptake (Fig. 1F) and lactate secretion (Fig. 1G), both of which were inhibited by Dex. Acute addition of LPS also caused an increase in glycolysis, which was detectable within 10-20 minutes (Fig. 1H). However this rapid response was insensitive to the presence of Dex. We hypothesise that the very rapid increase of glycolysis in response to acute LPS addition is mediated by proximal signaling events, whereas the long-term commitment to glycolysis requires changes of gene expression.

**Figure 1.**
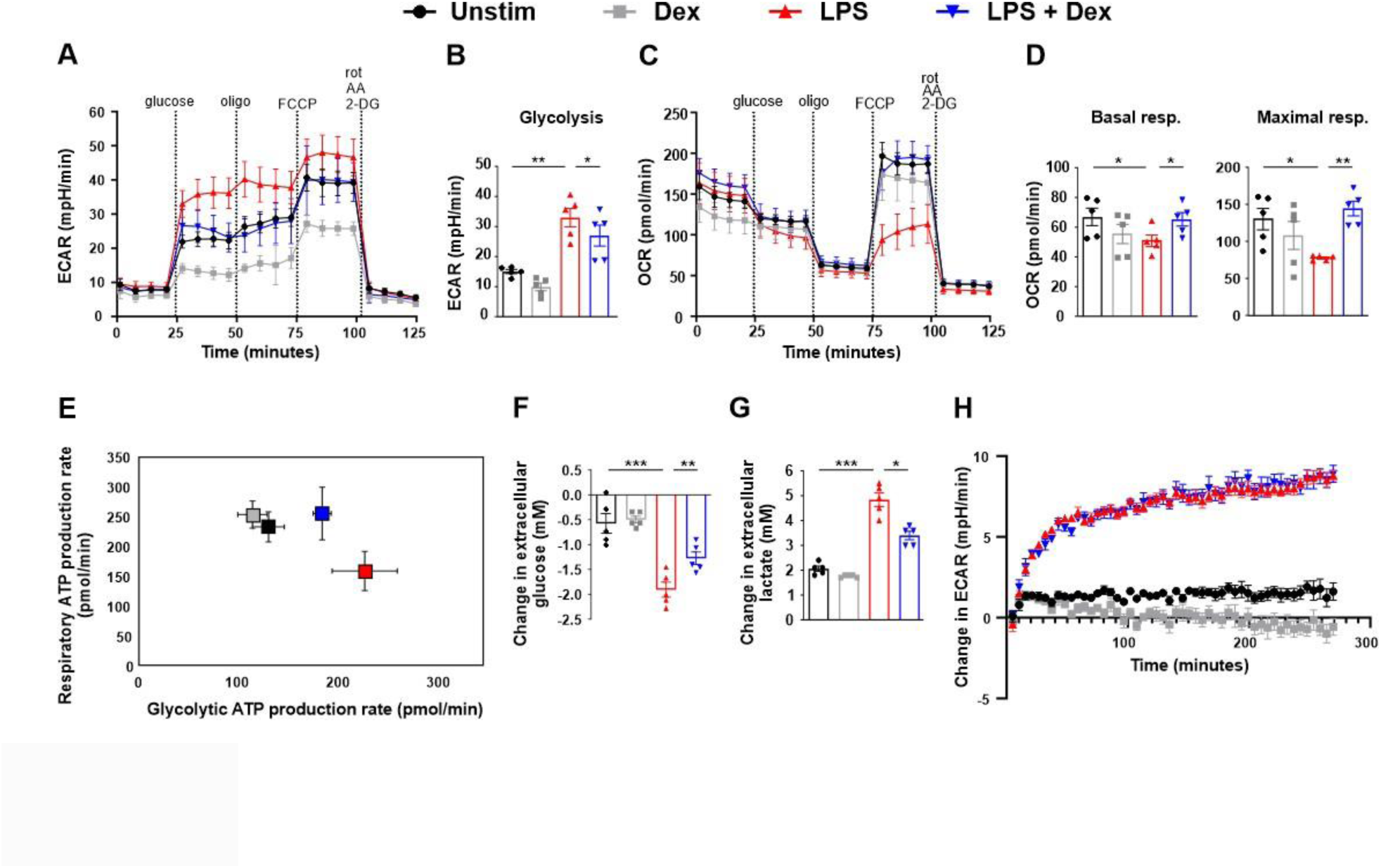
LPS-induced metabolic changes are opposed by dexamethasone in mouse bone marrow-derived macrophages. **(A-G)** Mouse BMDMs were treated for 24 h with vehicle (Unstim), LPS (10 ng/ml), Dex (100 nM) or a combination. **(A-D)** Mito+Glyco stress test performed using Seahorse XFe96 analyser. Representative plot (mean ± SD) and calculated metabolic parameters; mean ± SEM of 5 independent experiments. **E)** ATP rate assay; representative plot (mean ± SD) of 1 of 3 independent experiments. **F)** Change in extracellular glucose concentration measured in conditioned media following 24 h treatment; mean ± SEM of 5 independent experiments. **G)** Change in extracellular lactate concentration measured in conditioned media following 24 h treatment; mean ± SEM of 5 independent experiments. **H)** Extracellular acidification rate measured by Seahorse XFe96 analyser from BMDMs following acute injection of vehicle (Unstim), LPS (10 ng/ml), Dex (100 nM) or a combination at the beginning of the assay, expressed relative to starting value; mean ± SEM of 3 independent experiments. **(B,D,F,G)** One-way ANOVA with Dunnett’s multiple comparison correction. ECAR = extracellular acidification rate; OCR = oxygen consumption rate.

Nitric oxide inhibits enzymes of the electron transport chain and TCA cycle, contributing to the down-regulation of respiration in rat or mouse myeloid cells (26–28, 31). We hypothesised that Dex prevents LPS-induced impairment of respiration by inhibiting the expression of *Nos2* in mouse BMDMs. Indeed, Dex significantly inhibited LPS-induced expression of *Nos2* mRNA (Fig. 2A), and also had striking inhibitory effects on the expression of iNOS protein (Fig. 2B) and the production of nitric oxide (Fig. 2C). Inhibition of iNOS enzymatic activity can rescue oxidative phosphorylation in activated mouse myeloid cells (28). Dex may also preserve mitochondrial function in LPS-activated mouse BMDMs in part by preventing the accumulation of nitric oxide.

**Figure 2.**
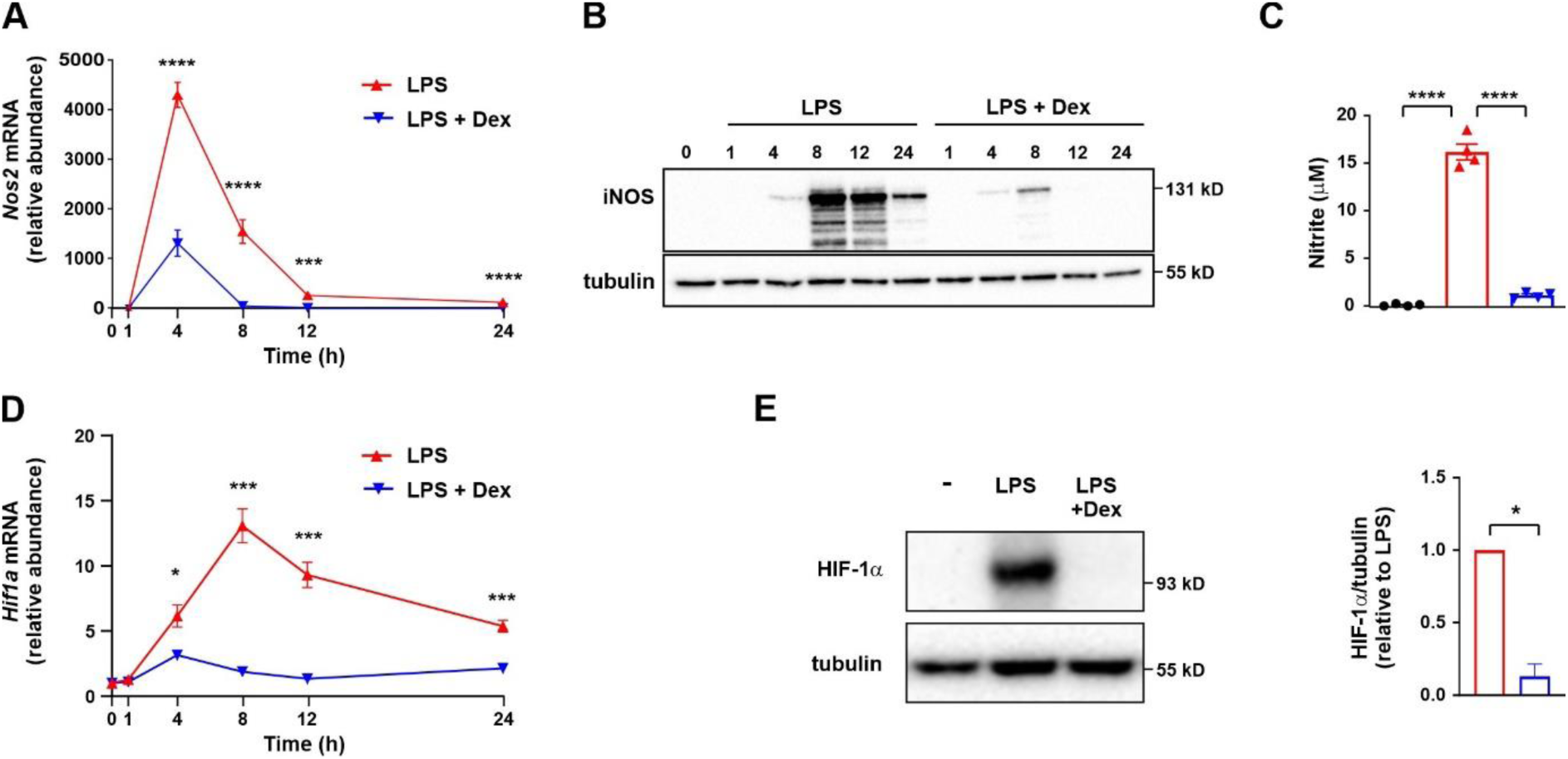
LPS-induced HIF-1*α* activation and *Nos2* expression are impaired by dexamethasone in mouse bone marrow-derived macrophages. Mouse BMDMs were treated for the indicated times with LPS (10 ng/ml) ± Dex (100 nM). **A)** *Nos2* mRNA was detected by RT-qPCR, normalised to housekeeper, expressed relative to unstimulated cells; mean ± SEM of 3 independent experiments; two-way ANOVA with Dunnett’s correction for multiple comparison. **B)** iNOS protein was detected by Western blotting. Representative image of 1 of 3 independent experiments. **C)** Nitrite was detected by Griess reaction on conditioned media following 24 h stimulation; mean ± SEM of 4 independent experiments; one-way ANOVA with Dunnett’s correction for multiple comparison. **D)** *Hif1a* mRNA expression was analysed by RT-qPCR, normalised to housekeeper, expressed relative to unstimulated cells; mean ± SEM of 3 independent experiments; two-way ANOVA with Dunnett’s correction for multiple comparison correction. **E)** HIF-1α protein was detected by Western blotting following 8 h stimulation. Representative image and quantification; mean ± SEM of 5 independent experiments.

### Dexamethasone inhibits HIF-1α activation and HIF-1α-dependent gene expression in LPS-treated mouse macrophages

LPS-induced expression of *Nos2* in mouse BMDMs is partly dependent on HIF-1α. (14, 28, 32). We therefore investigated the effect of Dex on LPS-induced HIF-1α activation. LPS-induced expression of *Hif1a* mRNA in mouse BMDMs was strongly inhibited by 100 nM Dex across a 24 hour time course (Fig. 2D). Measured at 8 h (at or near the peak of expression), LPS-induced expression of HIF- 1α protein was decreased on average 87% by Dex (Fig. 2E).

We hypothesised that Dex prevents LPS-induced glycolysis by impairing the activation of HIF-1α. To test this hypothesis we first validated a means of blocking HIF-1α function. The HIF-1α inhibitor KC7F2 (33) inhibited the hypoxic activation of an HRE-dependent reporter in HeLa cells (Fig. 3A). In LPS-treated BMDMs, KC7F2 impaired the accumulation of HIF-1α protein (Fig. 3B); the expression of two established HIF-1α targets, iNOS and pro-IL-1β (13, 14, 28, 29, 32) (Fig. 3C); and the activation-induced increase of glycolysis (Fig. 3D). We then used a published data set (14) to identify genes whose expression in LPS-treated BMDMs was dependent on HIF-1α (Fig. 3E). We selected a subset of highly HIF-1α dependent genes (*Slc2a1*, *Tpi1*, *Ldha* and *Slc16a3*; illustrated in Fig. 3F) and confirmed that their LPS-induced expression was inhibited by KC7F2 (Fig. 3G). There was a slight cytotoxic effect of KC7F2 at the higher dose of 40 µM, revealed by lower levels of α- tubulin protein in Fig. 3B and 3C. However, effects on glycolysis and glycolytic gene expression were also seen at the lower dose of 20 µM, at which no cytotoxicity was observed. Finally, we tested the prediction that the LPS-induced expression of HIF-1α-dependent genes would be significantly inhibited by Dex. In all cases this prediction proved to be correct (Fig. 3H). Notably, Dex prevented the LPS-induced expression of *Slc2a1*, encoding GLUT1, which is indispensable for LPS-induced glycolysis in BMDMs (34). Dex also significantly impaired the expression of the HIF-1α-dependent glycolytic genes *Hk2*, *Pdk1* and *Eno1* (not shown). Therefore Dex inhibits HIF-1α activation and expression of HIF-1α-dependent metabolic and pro-inflammatory genes in LPS-activated BMDMs.

**Figure 3.**
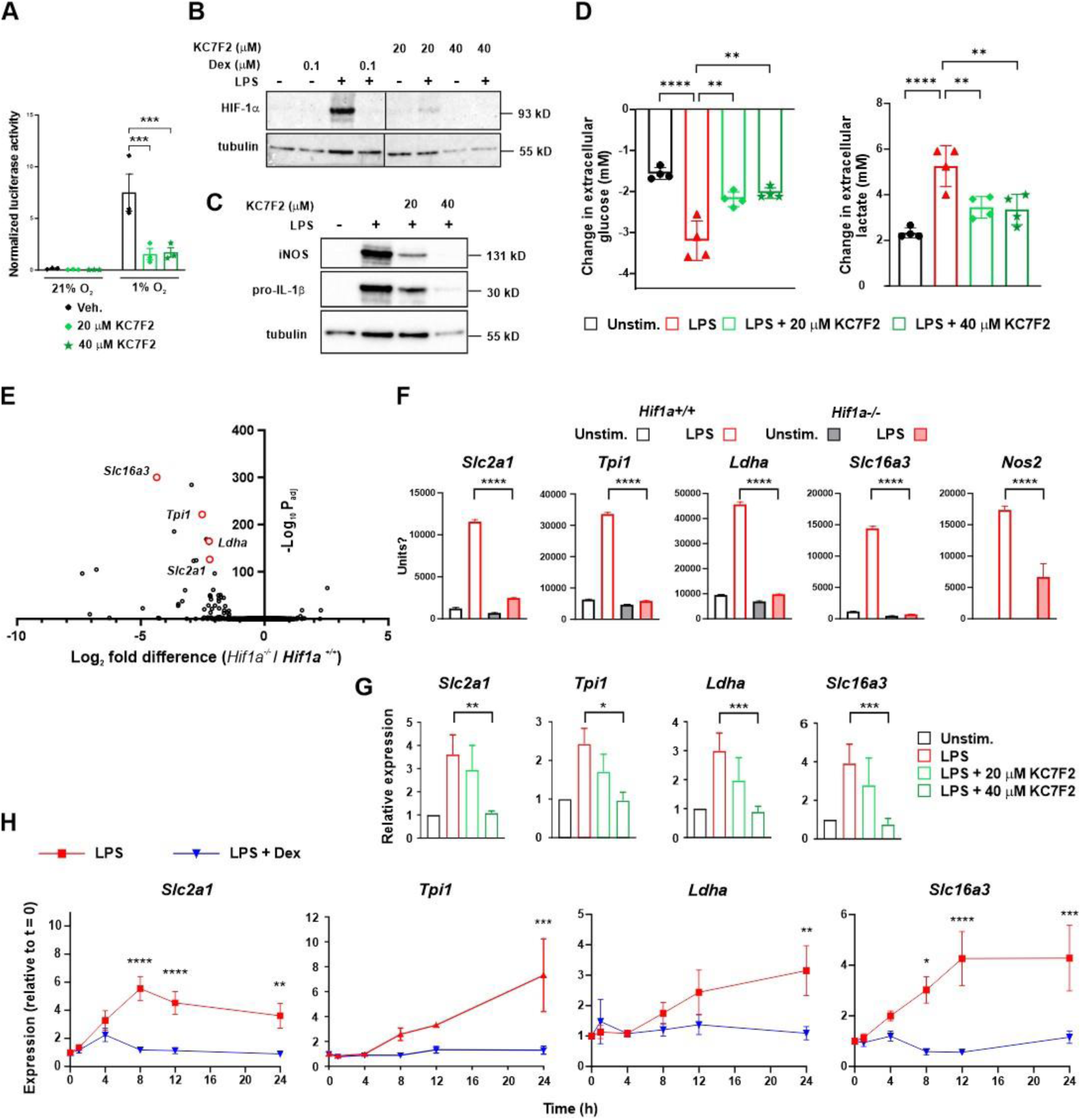
Dexamethasone impairs HIF-1*α*-mediated gene expression in LPS-treated bone marrow-derived macrophages. **A)** HRE-luciferase reporter assays were performed in HeLa cells. Cells transfected with reporter and control vectors were treated with vehicle (0.1% DMSO) or the indicated concentrations of HIF-1α inhibitor KC7F2 for 24 h, alongside incubation in normoxia or hypoxia (1% O_2_). Firefly luciferase signal was normalised to Renilla luciferase signal; mean ± SEM of 3 independent experiments; two-way ANOVA with Sidak’s correction for multiple comparison. **B,C)** Western blotting of whole cell lysates of BMDMs treated with LPS (10 ng/ml), Dex (100 nM), and the indicated concentrations of KC7F2 for 8 h. Representative images of 2 independent experiments. **D)** Change in extracellular glucose or lactate concentration measured in conditioned media following 24 h treatment; mean ± SEM of 4 independent experiments; one-way ANOVA with Dunnett’s correction for multiple comparison. **E)** Volcano plot of expression of LPS-induced genes in *Hif1a^+/+^*and *Hif1a^-/-^* BMDMs (our analysis of RNAseq data from GSE131364). **F)** Expression of selected genes in *Hif1a^+/+^* and *Hif1a^-/-^* BMDMs treated with vehicle (Unstim.) or LPS (100 ng/ml) for 18 h (our analysis of RNAseq data from GSE131364). **G)** Expression of HIF-1α-dependent genes was assessed by RT-qPCR from BMDMs treated with LPS (10 ng/ml) and the indicated concentrations of KC7F2 for 24 h. Expression was normalised against the untreated control (Unstim.); mean ± SEM of 4 independent experiments; one-way ANOVA with Dunnett’s correction for multiple comparison. **H)** BMDMs were treated for the indicated times with LPS (10 ng/ml) ± Dex (100 nM). Expression of genes of interest was quantified by RT-qPCR, normalised to housekeeper, expressed relative to unstimulated cells; mean ± SEM of 3-6 independent experiments; two-way ANOVA with Sidak’s correction for multiple comparison.

### Metabolic responses of primary human monocyte-derived macrophages to LPS and Dex differ from those of mouse bone marrow-derived macrophages

We then turned our attention to the metabolic responses of primary human monocyte-derived macrophages (MDMs). These cells also mounted a robust increase of glycolytic metabolism in response to LPS, and again this was significantly inhibited by Dex, as measured by extracellular acidification rate (Fig. 4A,B), glucose consumption (Fig. 4C) and lactate secretion (Fig. 4D). On the other hand, LPS did not cause any impairment of mitochondrial respiration in human MDMs; rather it caused a small but significant increase of basal respiration, which was further increased by addition of Dex (Fig. 4E,F). Maximal respiration was not significantly affected by either LPS or Dex (Fig. 4F). Differences from the metabolic responses of mouse BMDMs were also illustrated by ATP rate assays (Fig. 4G). LPS caused a right-ward but not down-ward shift indicating an increase in glycolytic ATP generation but no impairment of mitochondrial ATP generation. The LPS-induced increase in glycolytic ATP generation was inhibited by Dex. We and others have previously reported that LPS-activated human MDMs do not express *NOS2* or generate nitric oxide (19–22). This may help to explain why no LPS-induced impairment of mitochondrial function was detected.

**Figure 4.**
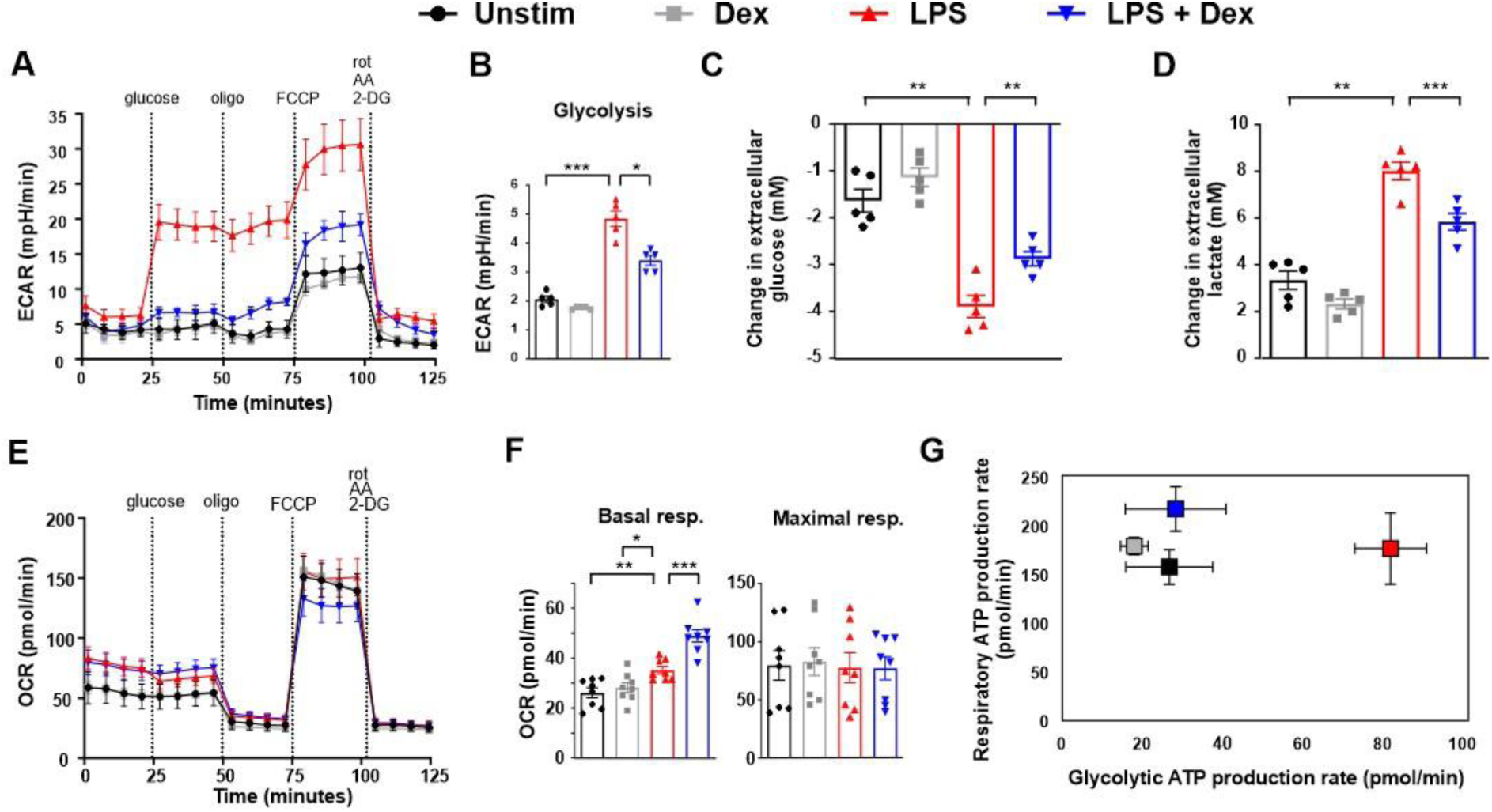
LPS-induced glycolysis is opposed by dexamethasone in human monocyte-derived macrophages. Human MDMs were treated for 24 h with vehicle (Unstim), LPS (10 ng/ml), Dex (100 nM) or a combination. Mito+Glyco stress tests were performed using Seahorse XFe96 analyser. **A)** Representative plot of extracellular acidification rate (ECAR); mean ± SD. **B)** Glycolysis rates were calculated from 8 independent experiments; mean ± SEM; one-way ANOVA with Dunnett’s correction for multiple comparison. Change in extracellular glucose **C)** or lactate **D)** concentration in conditioned media following 24 h treatment; mean ± SEM of 5 independent experiments; one-way ANOVA with Dunnett’s correction for multiple comparison. **E)** Representative plot of oxygen consumption rate (OCR) under conditions as described above. **F)** Basal and maximal respiration rates were calculated from 8 independent experiments; mean ± SEM; one way ANOVA with Dunnett’s correction for multiple comparison. **G)** ATP rate assay, representative plot (mean ± SD) of 7 independent experiments.

Because Dex impaired LPS-induced glycolysis in human MDMs as previously shown in mouse BMDMs, we investigated its effect on HIF-1α expression. Dex did not significantly inhibit LPS-induced expression of *HIF1A* mRNA (Fig. 5A), but consistently decreased the expression of HIF-1α protein (Fig. 5B). This suggested a post-transcriptional or post-translational mode of action. We therefore used cycloheximide chases to investigate effects of Dex on HIF-1α protein stability (Fig. 5C). In human MDMs treated for 8 h with LPS or LPS + Dex, HIF-1α protein decayed more rapidly in the latter. LPS-induced stabilisation of HIF-1α protein is not formally demonstrated here because HIF-1α protein could not be detected in the absence of LPS. However, the most parsimonious interpretation is that Dex reduces HIF-1α expression at least in part by interfering with LPS-induced protein stabilisation. Interestingly, the induction of HIF-1α protein by hypoxia, the combination of LPS and hypoxia (Fig. 5D) or the hypoxia mimetic dimethyloaxalylglycine (DMOG) (Fig. 5E) was relatively insensitive to Dex.

**Figure 5.**
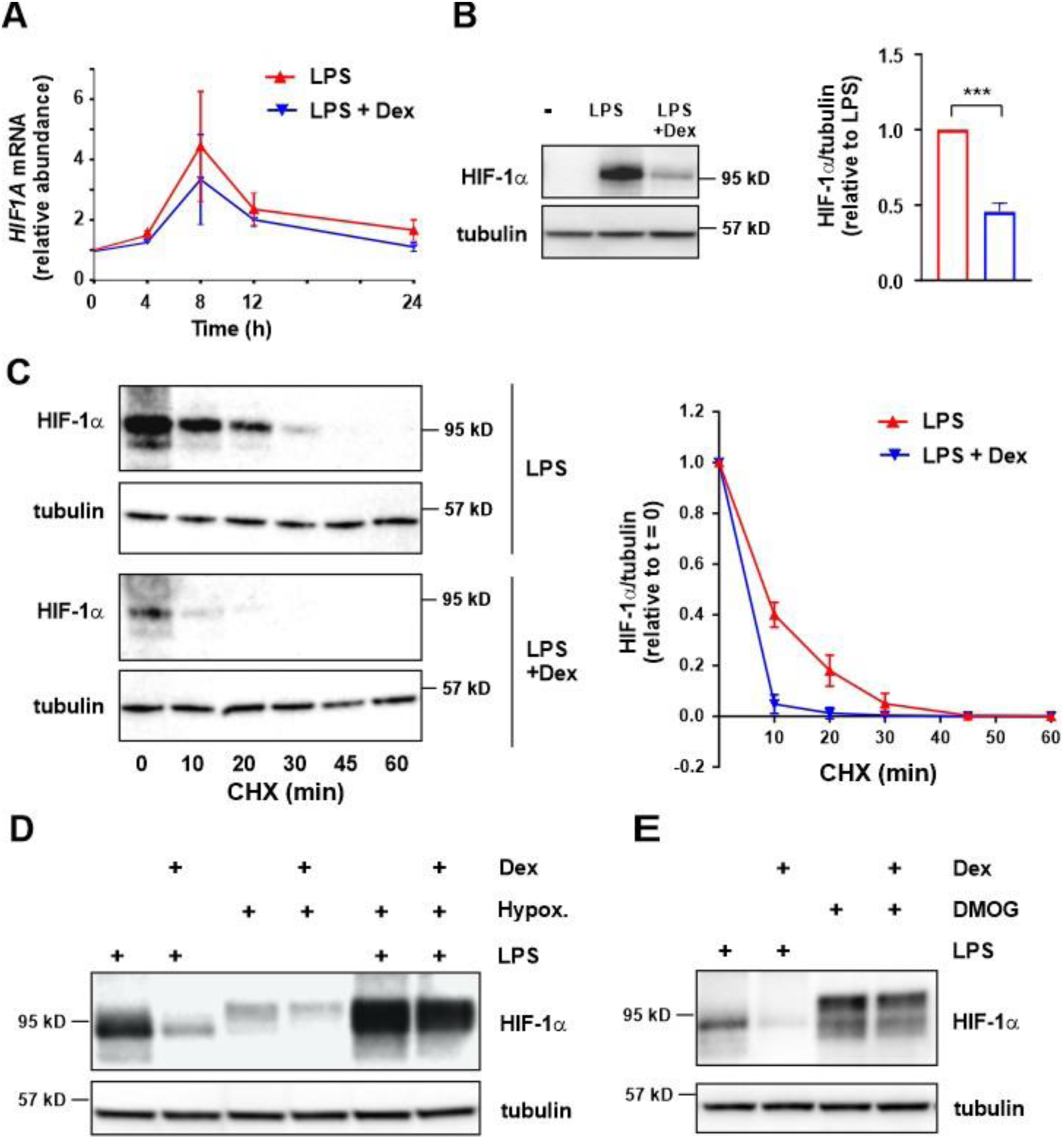
Dexamethasone destabilises HIF-1*α* and inhibits HIF-1*α* protein expression in LPS-treated human monocyte-derived macrophages. **A**) Human MDMs were treated with LPS (10 ng/ml) ± Dex (100 nM) for the indicated times. *HIF1A* mRNA was detected by RT-qPCR, normalised to housekeeper and expressed relative to unstimulated cells; mean ± SEM of 3 independent experiments. **B)** HIF-1α protein was detected by Western blotting following 8 h treatment of MDMs with LPS (10 ng/ml) ± Dex (100 nM). Representative image (left) and quantification of 10 independent experiments (right). **C)** Cycloheximide (CHX) chase and Western blot to measure HIF-1α protein stability. Following treatment with LPS (10 ng/ml) ± Dex (100 nM) for 8 h, cycloheximide (5 µg/ml) was added at t = 0 and cells were harvested after the indicated times. Representative image of 1 of 3 independent experiments (left); HIF-1α protein level (mean ± SEM), normalised against α- tubulin loading control and expressed relative to t = 0 (right). **D,E)** HIF-1α protein was detected by Western blotting of MDM lysates following 8 h treatment with LPS (10 ng/ml), dimethyloxallyl glycine (DMOG; 1 mM) or hypoxia (1% O_2_), all ± Dex (100 nM). Representative of at least 3 independent experiments for each condition.

We then proceeded to investigate the effects of LPS and Dex on glycolytic genes. Surprisingly, orthologues of most genes that were robustly up-regulated by LPS and suppressed by Dex in mouse BMDMs (Fig. 3H) were not responsive to either agonist in human MDMs (Fig. 6A) in spite of the high level of HIF-1α expression (Fig. 5B). Differences in expression of *TPI1* were statistically significant but rather small. In contrast the GLUT1-encoding transcript *SLC2A1* was strongly induced by LPS and inhibited by Dex. Western blotting (Fig. 6B) and flow cytometry (Fig. 6C) were used to confirm that GLUT1 protein expression was also increased by LPS and decreased by Dex. To establish that GLUT1 is also regulated by HIF-1α in human MDMs we first demonstrated that KC7F2 dose-dependently inhibited LPS-induced activation of HIF-1α in these cells, in this case without any evidence of cytotoxicity (Fig. 6D); then that 40 µM KC7F2 significantly inhibited the LPS-induced expression of *SLC2A1* mRNA (Fig. 6E). Therefore, in human MDMs, LPS activates HIF-1α and increases the expression of GLUT1 to support enhanced glycolysis, whereas Dex impairs HIF-1α activation, GLUT1 expression and LPS-induced glycolysis. Other effects of LPS and Dex on the glycolytic pathway have not yet been identified but cannot be formally ruled out.

**Figure 6.**
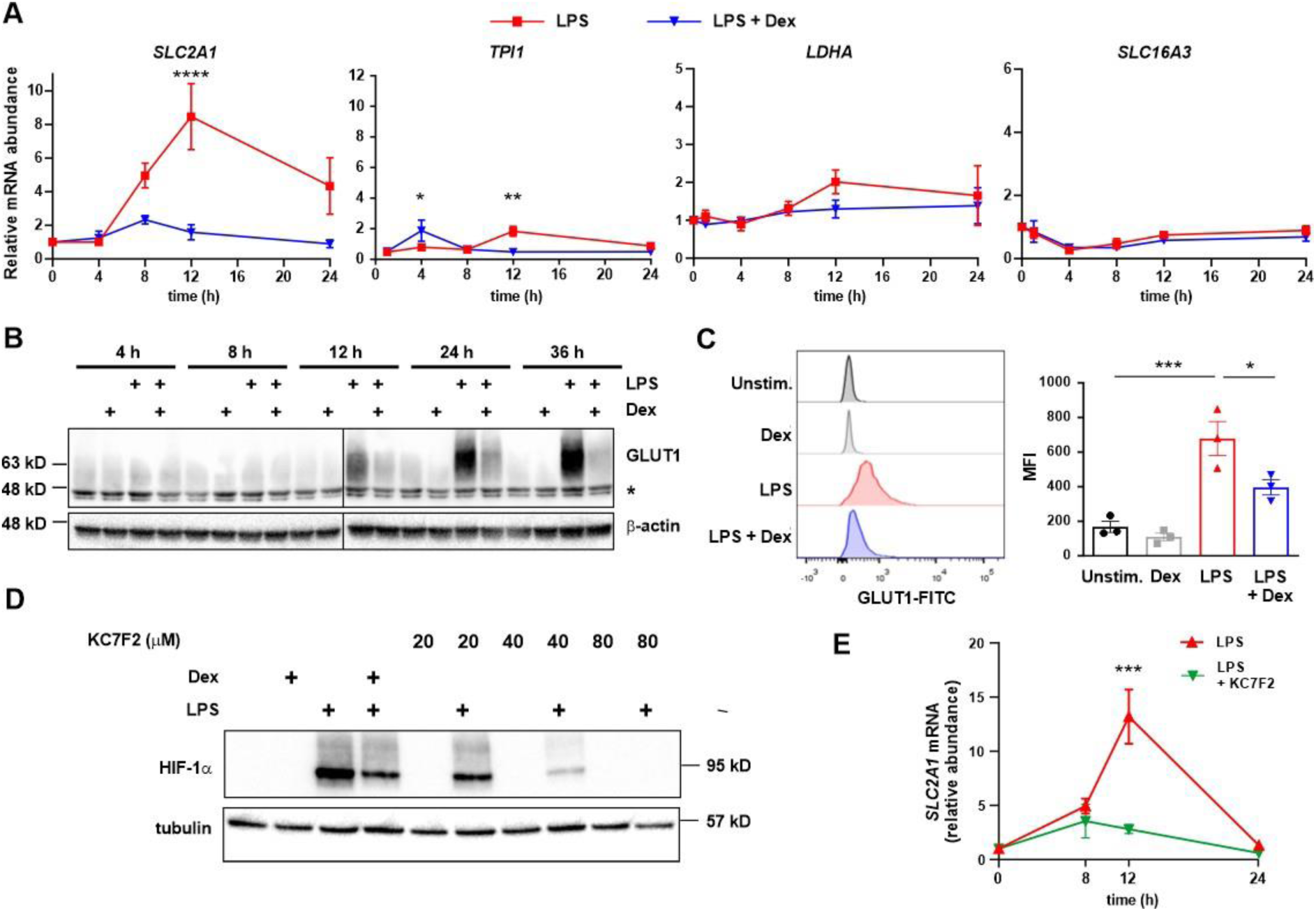
Dexamethasone inhibits expression of the HIF-1*α*-dependent gene *SLC2A1* in LPS-treated human monocyte-derived macrophages. **A)** Human MDMs were treated with LPS (10 ng/ml) ± Dex (100 nM) for the indicated times. Expression of selected glycolysis-related genes was measured by RT-qPCR and expressed relative to untreated controls; mean ± SEM of 3 independent experiments; two-way ANOVA with Dunnett’s correction for multiple comparison. To aid direct comparison, y-axes are the same as those used in Fig. 3H, where expression of the orthologous mouse genes was measured. **B)** GLUT1 protein was detected by Western blotting of whole cell lysates of MDMs treated with LPS (10 ng/ml) ± Dex (100 nM) as indicated. A representative of 3 independent experiments is shown. * indicates a non-specific band. **C)** GLUT1 protein was detected by flow cytometry of MDMs treated for 24 h with LPS (10 ng/ml) ± Dex (100 nM). Representative flow cytometry plots from 1 of 3 independent experiments are shown (left), and GLUT1 median fluorescent intensity (MFI) mean ± SEM plotted (right). **D)** MDMs were treated for 8 h with LPS (10 ng/ml) ± Dex (100 nM) or KC7F2 at the indicated concentrations. Whole cell lysates were blotted for HIF-1α. Representative of three independent experiments. **E)** MDMs were treated with LPS (10 ng/ml) ± KC7F2 (40 µM) for the indicated times. *SLC2A1* mRNA was detected by RT-qPCR, normalised to housekeeper and expressed relative to untreated controls; mean ± SEM of 3 independent experiments; two-way ANOVA with Dunnett’s correction for multiple comparison.

## DISCUSSION

GCs are stress hormones, whose immunosuppressive and anti-inflammatory effects have been known since the pioneering work of Philip Hench and others in the mid 20^th^ century. It seems paradoxical that under stressful conditions, even under existential threat, an organism will use endogenous GCs to effectively switch off a physiological system that could be essential for survival. We suggest that this paradox is best understood from an evolutionary perspective. Taking this approach, Ruslan Medzhitov and colleagues have elegantly described how metabolic decisions within immune cells reflect metabolic strategies at the level of the organism (35, 36). Under optimal conditions anabolic mechanisms support growth and reproduction, whereas in the presence of existential threats such as nutrient insufficiency, catabolic processes are favoured. The hypothalamus serves as a master regulator of these systemic switches by signaling through growth hormone and insulin-like growth factor, the hypothalamic-pituitary-gonadal (HPG) axis, or the hypothalamic-pituitary-adrenal (HPA) axis. As major consumers in the metabolic market place, immune cells are sensitive to the same hypothalamus-derived signals, and direct their metabolic activities in similar ways.

This evolutionary perspective should also explain how competing metabolic demands of different physiological systems are balanced and prioritised in such a way that the survival and reproductive success of the organism are optimised. For example, GCs produced in response to stress-induced activation of the HPA axis act on peripheral tissues such as muscle and white adipose to restrict uptake of glucose, such that it is conserved for the central nervous system (1, 37). Here we argue that the effects of GCs on immune cells might be understood in the same framework, their central function being to regulate glucose availability. In other words, GCs do not inhibit immune functions such as inflammation *in order to* conserve glucose for the brain; they restrict glucose utilisation by immune cells, and impair immunity and inflammation as an *unavoidable consequence* of the prime directive to maintain the supply of glucose to the brain. This hypothesis simplifies our understanding of the actions of both endogenous and synthetic GCs. If it is correct, it implies a commonality between the harmful effects of synthetic GCs, which are essentially metabolic in nature, and the desired anti-inflammatory effects, which are also based (at least partly) on alterations of metabolism. The extent to which harmful and beneficial effects of synthetic GR ligands can be separated then depends on whether GR regulates metabolism similarly in immune cells and metabolic tissues such as muscle and adipose. This is not yet known.

Review articles sometimes present descriptions of macrophage immunometabolism that are largely based on experiments in the mouse, with the implicit assumption that these are generalisable to other mammals. Here we add to a growing literature documenting metabolic differences between human and mouse macrophages (18–22). However, it is striking that broadly similar outcomes were generated by somewhat different means. In human MDMs, *HIF1A* expression was induced relatively weakly by LPS and not significantly affected by Dex. It is likely that the strong LPS-induced increase of HIF-1α levels in these cells is mediated largely by protein stabilisation, which is opposed by Dex. In mouse BMDMs, *Hif1a* mRNA was quite strongly induced by LPS, and this response was effectively blocked by Dex. Due to the extremely low levels of expression in the presence of Dex, we could not determine whether Dex also destabilised HIF-1α protein. In both mouse and human macrophages the outcome was similar: Dex-mediated suppression of LPS-induced HIF-1α protein expression. In mouse BMDMs this could explain the reduced expression of iNOS, pro-IL-1β and several genes of the glycolytic pathway, including the glucose and lactate transporters *Slc2a1* and *Slc16a3*. Although many glycolytic genes are well-established HIF-1α targets in other human cell types, (38–40), of the genes examined in human MDMs, only *SLC2A1* was positively and negatively regulated by LPS and Dex, respectively. We are currently investigating whether chromatin accessibility and HIF-1α recruitment at glycolytic genes differ between human and mouse macrophages. In any case, the effects of LPS and Dex on aerobic glycolysis were similar in macrophages of both species, whether this process was regulated via a single gene or several.

More striking differences were seen at the level of mitochondrial function. LPS treatment of mouse BMDMs caused impairment of mitochondrial respiration, which has previously been ascribed to reprogramming of mitochondria for the generation of reactive oxygen species, mediated (in part) by *Nos2* up-regulation and NO generation. *Nos2* expression and NO generation were strongly opposed by Dex. One implication of these findings is that exposure of murine macrophages to GCs could preserve their differentiation plasticity by preventing irreversible NO-mediated damage to the mitochondrial respiratory machinery, consequent commitment to aerobic glycolysis and pro-inflammatory functions (27, 28). On the other hand, the lack of *NOS2* expression and NO generation in activated human MDMs may help to explain why LPS did not cause impairment of mitochondrial oxidative phosphorylation in these cells, and suggests that these cells may retain the ability to re- polarise from M1-like (pro-inflammatory) to M2-like (reparative) phenotypes, rather than becoming committed to pro-inflammatory functions after exposure to LPS. We are currently testing these hypotheses.

It was notable that the LPS-induced expression of HIF-1α protein was highly sensitive to Dex, whereas Dex had relatively little effect on HIF-1α expression in response to hypoxia, a hypoxia mimetic or hypoxia plus LPS (Fig. 5). We also noted stimulus-dependent differences in electrophoretic mobility of HIF-1α, suggestive of different post translational modifications (Fig. 5). These observations raise some questions about mechanisms of activation of HIF-1α by LPS. According to current dogma, hypoxia, hypoxia mimetics and LPS all exert their effects by impairing prolyl hydroxylase activity, preventing hydroxylation and degradation of HIF-1α. Differential sensitivity to Dex suggests that there may in fact be differences in mechanism of activation. To give one hypothetical example, LPS could drive accumulation of HIF-1α by promoting a post-translational modification that impairs the recruitment of prolyl hydroxylase(s) (or by impairing a post-translational modification that favours prolyl hydroxylase recruitment). There are precedents for such indirect means of promoting HIF-1α accumulation (41–43), although none involving immune cells. In this case the target of Dex action could be a mediator of post-translational modification of HIF-1α rather than prolyl hydroxylase enzyme itself. Such a subtle difference of mechanism might explain why sensitivity to Dex varies according to stimulus. It will be important to answer these questions in order to identify ways to intervene in the metabolic reprogramming of macrophages in inflammatory contexts, and reduce inflammatory pathologies that are driven by macrophages.

GCs have been previously reported to influence HIF-1α function in various leukocyte populations. High doses of GCs are used in the treatment of lymphocytic malignancies such as acute lymphoblastic leukaemia, where their pro-apoptotic effects have been linked to down-regulation of GLUT1 expression, inhibition of glucose consumption and glycolytic flux (44–46). Dexamethasone was shown to inhibit hypoxia-induced expression of HIF-1α protein in CD4+ T cells (47). Intravenous treatment of multiple sclerosis patients with the GC methylprednisolone decreased the expression in peripheral T cells of metabolic genes, including *LDHA* (48). In myeloid-derived suppressor cells dexamethasone inhibited the expression of HIF-1α protein and certain glycolytic genes (*Slc2a1*, *Eno1* and *Slc16a3*) (49). Finally, a recent publication reported that dexamethasone inhibited HIF-1α protein accumulation and glycolytic metabolism in LPS-treated mouse BMDMs (50), as described here. However, in that study GC-mediated changes in expression of HIF-1α-dependent glycolytic genes was not reported.

In summary, we have shown here that the synthetic GC dexamethasone acts on both mouse and human macrophages to inhibit LPS-induced activation of HIF-1α, and to oppose the metabolic reprogramming that is dependent on HIF-1α. There may be parallel effects of GCs on the HIF-1α pathway in other immune cells. Our findings may have important implications for how we understand beneficial and harmful effects of GCs, and whether it will be possible to separate these in a meaningful way.

## DATA AVAILABILITY

We downloaded and analysed raw RNAseq data from a publicly accessible database (GSE131364). All other data underlying this article are included in the manuscript. Raw images or data files will be made available upon reasonable request to the corresponding author.

## ETHICS APPROVAL

Human monocyte-derived macrophages were generated from anonymous donor leukapheresis cones under ethical approval ERN_16-0191. Mouse colonies were maintained according to the Home Office guidelines and approved by the University of Birmingham Animal Welfare and Ethical Review Board.

## PATIENT CONSENT

Not applicable.

## PERMISSION TO REPRODUCE MATERIAL FROM OTHER SOURCES

Not applicable

## CLINICAL TRIAL REGISTRATION

Not applicable.

## ACKNOWLEDGEMENTS

We are grateful to Kim and colleagues for deposition of RNAseq in the Gene Expression Omnibus (14), and to members of the Clark and Tennant groups for helpful discussions. Graphical abstract was produced using BioRender.

